# Autonomic and cortical responses to heartbeat-synchronous auditory omissions during an auditory attention task

**DOI:** 10.64898/2026.07.14.738383

**Authors:** Mai Sakuragi, Yuto Tanaka, Kazushi Shinagawa, Yuri Terasawa, Satoshi Umeda

## Abstract

Cardiovascular regulation depends on bidirectional communication between autonomic cardiac control and cardiac afferent signals ascending to the central nervous system, forming a heart–brain loop. Cardio–auditory omission paradigms provide a non-invasive way to probe this loop in humans by withholding scheduled tones within heartbeat-contingent auditory sequences and measuring the resulting autonomic cardiac and heartbeat-related cortical responses. Previous studies have shown omission-evoked cardiac deceleration and neural responses mainly in specific contexts such as passive listening or synchrony-judgement tasks. The present study examined whether these responses also occur under the more general condition of active responses to external stimuli. We tested 40 adults in an auditory detection task in which tones were presented either 200 ms after the ECG R-peak (synch) or at pseudo-random intervals matched to each participant’s resting heart rate (asynch). A five-minute no-stimulation resting recording was used to generate the asynch sequence and served as the resting comparison condition. In both auditory conditions, 10% of scheduled tones were omitted, yielding synch and asynch omissions. Heartbeat-synchronous omissions induced sustained RR-interval prolongation that exceeded responses to asynch omissions, sound-present conditions, and rest. HEP amplitude was selectively enhanced in the synch-omission condition relative to all other conditions. Reaction times to the first post-omission tone were slightly delayed, whereas overall performance remained comparable across conditions. These findings show that heartbeat-synchronous auditory omissions produce a distinct autonomic–cortical response profile under active response demands, indicating that heart–brain loop dynamics continue to shape physiological responses during ongoing behaviour.

## 1. Introduction

The autonomic nervous system, and in particular cardiovascular control, plays a central role in how the brain coordinates bodily states with perception, emotion, and behaviour. Cardiac activity is not only regulated by central autonomic pathways but also continuously feeds back to the brain via vagal and glossopharyngeal afferent pathways, forming a bidirectional heart–brain loop embedded in central autonomic and viscerosensory networks (Critchley & Harrison, 2013; Park & Blanke, 2019). Within this loop, heartbeat-locked neural responses provide a complementary index of cortical responses to cardiac afferent signals. In particular, the heartbeat-evoked potential (HEP), obtained by averaging scalp or intracranial EEG time-locked to the ECG R-peak, has been used to index cortical responses associated with cardiac and baroreceptor-related inputs, although its physiological and functional interpretation remains complex. HEP amplitude has been reported to vary with several cardiovascular and task-related factors, including the behavioural relevance of bodily signals and the allocation of attention (Kern et al., 2013; Park & Blanke, 2019; Schulz et al., 2025). Cardiac and baroreceptor-related afferent signals ascend primarily via vagal and glossopharyngeal pathways to the nucleus tractus solitarius and associated brainstem relays, from which they reach thalamic, insular, cingulate, and somatosensory regions that are thought to contribute to HEP generation and autonomic reflex control (Berntson & Khalsa, 2021; Ferraro et al., 2022). Taken together, RR-interval dynamics and HEP amplitude provide complementary indices of the heart–brain loop, reflecting autonomic cardiac modulation and heartbeat-related cortical processing, respectively. Examining these measures together can therefore help clarify how cardiac control and cortical heartbeat responses are jointly expressed during ongoing behaviour.

Recent work using cardio–auditory omission paradigms has begun to characterise neural and cardiac responses to omitted tones within heartbeat-contingent auditory sequences. In these paradigms, tones are presented either at a fixed delay after the R-peak or in comparison sequences without such heartbeat locking, and occasional omissions allow responses to the absence of a scheduled tone to be measured under controlled timing conditions. In passive listening designs, omissions within regular heartbeat-synchronous tone trains elicit enhanced omission-locked potentials and a short-latency cardiac deceleration compared with omissions in irregular or asynchronous sequences (Pfeiffer & De Lucia, 2017). In synchrony-judgement tasks, the magnitude of omission-evoked HEP modulation depends on the perceived cardio–audio delay and on whether the sequence is judged as synchronous or asynchronous. These findings suggest that cortical responses to omissions are sensitive to the temporal relationship between heartbeats and sounds (Banellis & Cruse, 2020). More recently, cardio–audio synchronisation has been shown to elicit omission-locked neural responses and heartbeat deceleration during wakefulness, non-REM sleep, and coma after cardiac arrest (Pelentritou et al., 2024, 2025). Together, these studies show that cardio–auditory omissions can evoke neural and cardiac responses across several specific contexts, including passive listening, synchrony judgement, and reduced-consciousness states.

However, it remains unclear whether these coupled autonomic and cortical responses are also expressed under the more general condition of active responses to external stimuli. Passive listening to heartbeat-synchronous sounds and explicit synchrony judgements provide important experimental control, but they differ from ordinary behavioural contexts in which external events must be monitored and acted upon. This issue is physiologically relevant because cardiovascular control and cortical processing of cardiac signals do not occur in isolation during ongoing behaviour, but operate while external events are processed and behavioural responses are prepared. If the cardiac deceleration and neural responses observed in previous cardio–auditory omission studies (Banellis & Cruse, 2020; Pelentritou et al., 2024, 2025; Pfeiffer & De Lucia, 2017) also occur under such response demands, this would indicate that heart–brain loop dynamics remain expressed during ongoing stimulus-guided behaviour, rather than being limited to passive listening or explicit synchrony-judgement contexts.

The present study examined autonomic cardiac modulation and heartbeat-related cortical responses to heartbeat-synchronous auditory omissions during an auditory detection task. Participants pressed a key whenever they heard a tone, while tones were presented either 200 ms after the ECG R-peak (synch condition) or at pseudo-random intervals matched to each participant’s resting heart rate (asynch condition). In both auditory conditions, 10% of scheduled tones were omitted, yielding synch and asynch omissions. A five-minute no-stimulation resting recording was used both to generate the asynch sequence and to provide a resting comparison condition. To determine whether the response to heartbeat-synchronous omissions was distinct from more general effects of sound omission, sound timing, or task engagement, we compared RR-interval dynamics and HEP amplitude across synch omissions, asynch omissions, sound-present conditions, and rest.

Based on previous findings that heartbeat-synchronous omissions evoke cardiac deceleration and enhanced omission-related or heartbeat-related neural responses (Banellis & Cruse, 2020; Pelentritou et al., 2024, 2025; Pfeiffer & De Lucia, 2017), we tested two primary hypotheses. First, at the autonomic level, we hypothesised that heartbeat-synchronous omissions would induce larger and more sustained RR-interval prolongation, indicating cardiac deceleration, than asynch omissions, sound-present conditions, and rest. Second, at the cortical level, we hypothesised that HEP amplitude would be selectively enhanced in the synch-omission condition relative to the other conditions. For the HEP analysis, heartbeat-locked epochs occurring after an omission and before the next tone were used for the omission conditions, and a subtraction procedure was applied to reduce auditory-evoked contributions.

We also conducted exploratory analyses of behavioural effects, attenuation of physiological responses across repeated omissions, and interoceptive individual differences. At the behavioural level, we examined whether omissions produced a transient delay in responses to the first post-omission tone and whether overall task performance remained preserved. At the physiological level, we examined whether omission-evoked RR and HEP effects showed clear attenuation across repeated trials. Finally, interoception refers to the sensing and evaluation of internal bodily signals and is commonly distinguished into dimensions such as interoceptive accuracy, sensibility, and awareness (Garfinkel et al., 2015; Khalsa et al., 2018). In the cardiac domain, heartbeat-perception tasks have been widely used as behavioural indices related to cardiac interoception (Schandry, 1981), whereas self-report questionnaires assess subjective bodily awareness or beliefs (Cabrera et al., 2018; Mehling et al., 2012). Although some studies have reported associations between HEP amplitude and measures related to cardiac interoception, the interpretation of these associations remains debated and may depend on the task and measurement approach (Coll et al., 2021; Pollatos & Schandry, 2004). In addition, previous cardio–auditory work suggests that individual differences in cardiac interoceptive measures may be relevant when participants explicitly attend to heartbeat–sound synchrony (Banellis & Cruse, 2020). We therefore explored whether there was any evidence that omission-evoked RR and HEP responses varied with interoceptive individual-difference measures in the present active auditory detection task.

## 2. Methods

### 2.1. Participants

Forty-two healthy undergraduate students, graduate students, and working adults participated in the experiment (19 males, 23 females; mean age = 22.2 years; age range = 18–30 years). All participants had normal or corrected-to-normal vision. The sample size was determined a priori using Bayes factor design analysis (BFDA) with the Sample Size Planner. In the BFDA, we assumed an analysis comparing heart-rate changes immediately after sound omissions across experimental conditions. Heart-rate change was defined as the percentage change in RR interval from RR-1 as the baseline to RR0–RR5. A repeated-measures ANOVA-based model, approximating a general linear mixed model, with Stimulus Condition (Synch/Asynch/Rest) and Epoch (RR0–RR5) was specified as the main design framework. The interaction effect, reflecting whether the temporal profile of heart-rate changes differed across conditions, was defined as the primary effect of interest. The required sample size was calculated with a true positive rate of 0.8, an expected standardized effect size of δ = 0.5, a Bayes factor threshold of BF10 = 3, and a prior distribution scale parameter of 1/√2. The stopping rule was fixed-N sampling with a maximum sample size, and the required sample size was set to N = 42 based on these settings. However, two participants were excluded from the analysis because of excessive noise in the electrocardiogram (ECG) recordings; therefore, the final analytic sample consisted of 40 participants.

Before participation, we screened participants for factors that could affect physiological recordings, such as EEG and ECG, or performance on the psychological task. Participants were required not to meet any of the following exclusion criteria: history of epileptic seizures; history of surgical procedures involving the head; cardiac disease; current use of medications affecting psychiatric or neurological functions, headache analgesics, cold medicine, or anti-allergic agents. Individuals who had difficulty understanding the experimental instructions or lecture content in Japanese, or who had consumed vasoactive substances, including food or drinks containing caffeine, alcohol, or nicotine, within three hours before the experiment were also excluded. Participants wearing makeup, particularly around the eyes, were asked to report this to the experimenter on the day of the experiment and to remove it if necessary. If they reported that they were unable to comply with this requirement, they were not allowed to participate. Whether participants met any of these criteria was confirmed verbally on the day of the experiment. No participants were excluded after applying to the study based on these criteria. All participants provided written informed consent before participation. The study was approved by the Keio University Research Ethics Committee (Approval No. 240040000; approved 7 May 2024) and was conducted in accordance with the Declaration of Helsinki.

### 2.2. Apparatus

Throughout all experimental tasks, ECG was measured using an MP-150 system and AcqKnowledge (Biopac Systems, Santa Barbara, CA), and continuous blood pressure was monitored using VitalStream (Caretaker Medical, Charlottesville, VA). ECG was recorded in a three-lead configuration with electrodes placed on the right hand and both feet, while continuous blood pressure was measured with a cuff on the right middle finger. During the attention task using heartbeat-synchronous sounds, EEG was recorded alongside ECG using NetStation 5.3.0.1 and a 64-channel HydroCel Geodesic Sensor Net. Respiration (changes in chest and abdominal circumference) was measured using a respiratory effort belt (RESP100C, Biopac Systems, Santa Barbara, CA), and pupil diameter during the task was recorded with an eye tracker (Tobii Pro Spectrum, Tobii, Stockholm, Sweden). In the present paper, however, we report only the ECG- and EEG-based results.

### 2.3. Questionnaires

Interoception-related self-report measures included the Body Awareness scale of the Body Perception Questionnaire (BPQ-BA; Kobayashi et al., 2021) and the Multidimensional Assessment of Interoceptive Awareness, Japanese version (MAIA-J; Shoji et al., 2018). These questionnaires were completed at home after the experimental session. BPQ-BA total scores were calculated by averaging item scores after reverse scoring where appropriate. For the MAIA-J, average scores were calculated for each subscale: Noticing, Not-Distracting, Not-Worrying, Attention Regulation, Emotional Awareness, Self-Regulation, Body Listening, and Trusting.

### 2.4. Procedure

#### 2.4.1. Heartbeat counting task (HCT)

To obtain a behavioural index related to cardiac interoception, participants completed the Heartbeat Counting Task (HCT; Schandry, 1981). The task included resting pulse measurement, heartbeat counting trials, and a Time Estimation Task (TET). During these tasks, participants placed their left arm on the table to enter responses, while their right hand, fitted with the continuous blood pressure monitor, remained at rest on their lap. In the HCT, participants silently counted their heartbeats during predefined intervals consisting of two trials each of 25, 35, and 45 s, and reported the count for each interval. Participants were instructed not to count by touching their body, pressing firmly against the backrest or desk, or holding their breath. Following the revised HCT instructions (Desmedt et al., 2018), they were also instructed not to guess heartbeats they could not feel and not to calculate the count based on prior knowledge of their heart rate or perceived elapsed time. After each HCT trial, participants rated their confidence in their performance on a scale from 1 to 10. The TET was then administered to assess individual differences in time estimation. Participants estimated predefined intervals consisting of two trials each of 23, 49, and 56 s and reported the perceived duration. The intervals for both HCT and TET were presented in a randomized order.

#### 2.4.2. Heartbeat-synchronous auditory task

Following the HCT, participants completed the STAI-State questionnaire to record their mood before the heartbeat-synchronous auditory task. Subsequently, in addition to the ECG and continuous blood pressure monitor, the EEG net and two respiratory belts (chest and abdomen) were fitted. Participants were seated comfortably in a sound-attenuated laboratory and instructed to remain still with their eyes open, right hand on their lap, and feet flat on the floor to ensure high signal quality.

The auditory stimuli consisted of pure tones (1000 Hz, 65 dB) with a duration of 100 ms. The stimulus conditions comprised a no-sound baseline and two auditory conditions: a heartbeat-synchronous condition (synch), in which tones were presented in temporal alignment with the heartbeat, and an asynchronous condition (asynch), in which tones were presented without such alignment. Details of the timing procedure are described below. The two auditory conditions were presented in separate experimental blocks, with the order randomized across participants. Participants were instructed to press the “9” key with their left hand as quickly and accurately as possible whenever they heard a sound. To acquire data for thought-related analyses in a separate project, thought probes assessing participants’ ongoing thought states were presented after every 50 to 70 key presses, with the exact interval randomly determined from a uniform distribution. Each trial was defined as ending when a thought probe was presented, and the task consisted of 15 trials in total. Participants completed the task in 25 to 30 minutes.

Before starting the task, five minutes of resting physiological data were acquired without auditory stimulation; this resting period served both to generate the sound sequence for the asynch condition and as the no-sound baseline condition. During this resting period, ECG was recorded and later analyzed offline. RR intervals were calculated using automatic R-peak detection, and the minimum and maximum RR intervals during the rest period were computed using R (ver. 4.4.1). In the experimental control software Presentation (Neurobehavioral Systems, Berkeley, CA), a pseudo-random sound sequence approximating each participant’s heart rate rhythm was then generated from a uniform distribution based on these minimum and maximum values. This procedure minimized differences in the mean and variance of the RR intervals between the synch and asynch conditions. After the resting measurements and generation of the asynch sequence, the auditory attention task was started.

The procedure for each condition is summarized in Figure 1. In both synch and asynch conditions, sound omissions occurred with a 10% probability. To prevent consecutive omissions, at least three sound stimuli were always inserted between omissions. In the synch condition, the onset of the auditory stimuli was triggered by R-peaks detected online from the raw ECG recording. To enable real-time R-peak detection, the raw ECG was analyzed in real time using AcqKnowledge (Biopac Systems, Santa Barbara, CA). An R-peak was detected when the online ECG value fell within the range of 0.3 to 0.4 μV, and the sound stimulus or omission was triggered 200 ms after detection. Since R-wave amplitudes varied among individuals, the detection range was adjusted in 0.1 μV increments to match each participant’s signal. The 200 ms delay was chosen based on the average delay associated with human perception of the heartbeat (Brener & Ring, 2016). In the asynch condition, the timing of stimulus presentation followed the pseudo-random sequence of RR intervals derived from the pre-recorded resting ECG, such that tones and omissions approximated the participant’s heart rate rhythm without being locked to the ongoing R-peaks.

**Figure 1.**
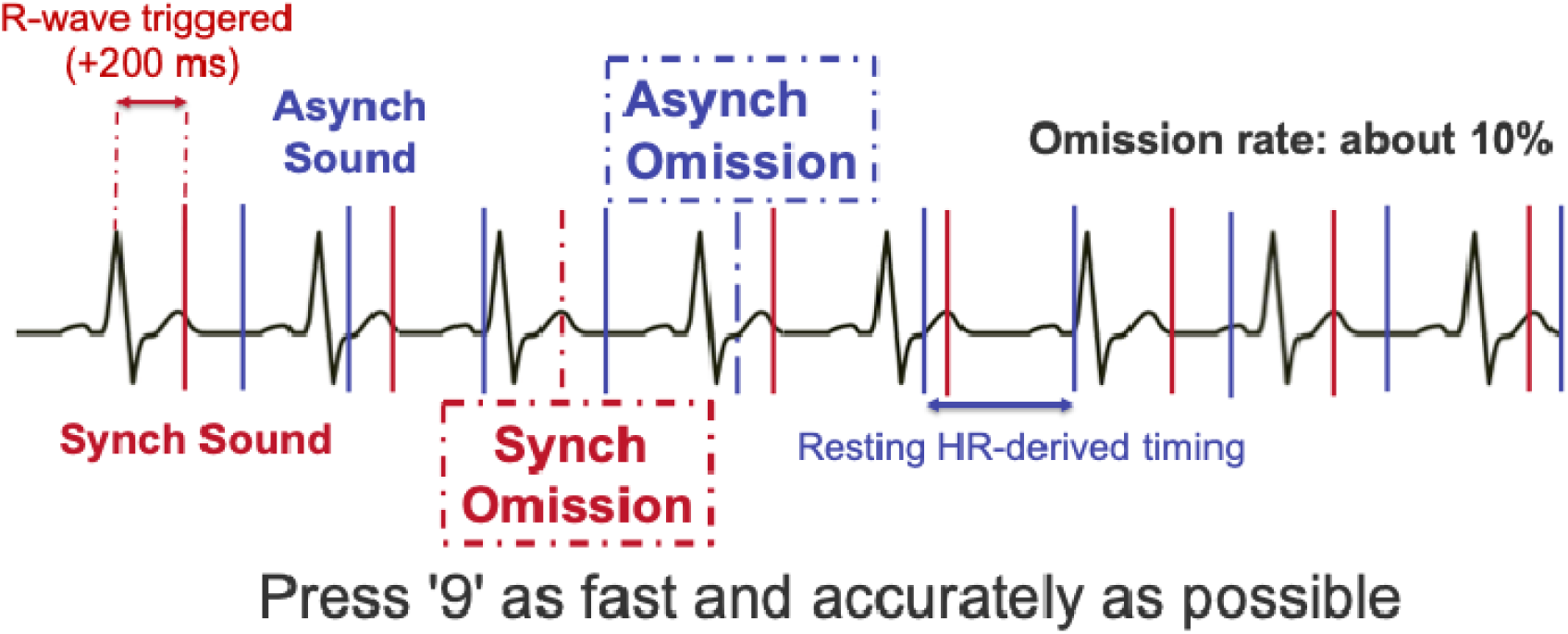
Experimental task procedure. Participants performed a task in which they were required to press the “9” key whenever an auditory stimulus was presented. In the synch condition, auditory stimuli were presented 200 ms after the detection of real-time R-peaks. In the asynch condition, stimuli were presented at intervals drawn from a uniform distribution between the minimum and maximum RR intervals extracted from a resting baseline block. Each condition consisted of 15 trials, with each trial defined as 50–70 key presses.

At the end of the experiment, participants were informed that auditory events had been presented according to different timing rules across blocks: one block followed a pseudo-random sequence based on resting heart rate, whereas the other was triggered by real-time R-peaks. Before this disclosure, participants were asked whether they had noticed any systematic difference in the timing of auditory events between blocks. No participant reported noticing this distinction before disclosure.

### 2.5. Data Preprocessing

#### 2.5.1. ECG data

R-peaks were detected from the raw ECG waveforms using AcqKnowledge, and RR intervals were calculated as the difference between adjacent R-peak timestamps. To exclude physiological artifacts, threshold processing was applied to each participant’s RR intervals using their median RR interval. Epochs exceeding 1.5 times the median RR interval or falling below 2/3 of the median RR interval were defined as outliers and excluded from the analysis. Physiological data were synchronized with the HCT and auditory attention task trials based on event triggers from the experimental control logs. To control for the effects of motor responses and transient cognitive load associated with responding to thought probes, the intervals from probe presentation to response completion were identified and excluded, extracting only the RR data during task performance.

#### 2.5.2. EEG data

Preprocessing of EEG data was performed using the EEGLAB toolbox (Delorme & Makeig, 2004) and its MATLAB (MathWorks, USA) plugins. Raw EEG data were downsampled to 250 Hz, and a band-pass filter was applied from 0.5 Hz to 40 Hz. Artifact Subspace Reconstruction (ASR) was used to remove and correct artifacts. First, bad channels were detected (correlation with adjacent channels < 0.85; high-frequency noise *SD* > 4) and interpolated using spherical splines. Subsequently, ASR correction was applied to intervals containing short-duration burst noise (Burst Criterion = 30 *SD*). After ASR, the data were re-referenced to the common average. Independent component analysis using the extended Infomax algorithm was performed to remove biological artifacts (eye movements, muscle activity, ECG). ICLabel (Pion-Tonachini et al., 2019) was used to automatically classify independent components. Components exceeding the following probability thresholds were excluded: Eye movements > 0.85, Muscle activity > 0.85, ECG > 0.60, Line noise > 0.85, and Channel noise > 0.85. Finally, a second strict cleaning (Burst Criterion = 15 *SD*, Window Criterion = 0.2) was performed to remove remaining significant artifacts.

#### 2.5.3. Questionnaire scores

Total scores for the STAI, BDI, and BPQ were calculated by dividing the sum of all items (considering reverse-coded items) by the number of items. For the MAIA-J, average values were calculated for each subscale: Noticing, Not-Distracting, Not-Worrying, Attention Regulation, Emotional Awareness, Self-Regulation, Body Listening, and Trusting.

### 2.6. Statistical Analysis

#### 2.6.1. Analysis of RR interval fluctuations associated with sound omissions

To examine heart rate changes immediately following sound omissions, the RR intervals surrounding each omission were identified: the interval immediately preceding the omission (RR-1), the interval containing the omission (RR0), and the five subsequent intervals (RR1 to RR5). The rate of change for RR0–RR5 was calculated relative to the RR-1 value. For comparison, a bootstrap sample of the same time window was extracted from the resting heart rate data. A fixed block containing seven data points (RR-1 to RR5) was sampled 1,000 times. A Bayesian statistical model was constructed to compare heart rate changes between the rest, synch, and asynch conditions. The dependent variable was the change in RR interval (RR-1 to RR5). Fixed effects included the main effects of Epoch (RR-1 to RR5) and Stimulus Condition, along with their interaction. Participants were included as a random effect.

Furthermore, to examine whether the heart rate change effect attenuated as the task progressed, a Bayesian model was constructed with the mean heart rate change per trial as the dependent variable, including the main impacts of Epoch and Trial Number, and their interaction. Both models assumed a normal distribution for the dependent variable. Prior distributions were set as follows: *Normal* (0, 1) for fixed effects and *Student-t* (*df* = 3, μ = 0, σ = 10) for the standard deviation of random effects. Sampling was performed with 1,000 warm-up iterations and 4,000 total iterations (4 chains).

#### 2.6.2. Calculation of heartbeat-evoked potentials (HEP) following sound omissions

EEG data were averaged using the ECG R-peak as the trigger. The analysis window was set from -100 ms to 500 ms relative to the R-peak. To prevent the cardiac field artifact (CFA) from contaminating the baseline, the average potential from -100 ms to -50 ms—a stable period immediately preceding the R-peak—was used as the baseline and subtracted from each epoch. Heartbeat epochs occurring during the interval from an omitted auditory stimulus to the subsequent auditory stimulus were averaged as HEPs for the synch/asynch omission conditions. In contrast, heartbeat epochs occurring during intervals in which auditory stimuli were actually presented were averaged as HEPs for the synch/asynch sound conditions.

In the synch sound condition, since auditory stimuli are presented in synchrony with the R-peak, the observed potentials inevitably include a superposition of the HEP and the Auditory Evoked Potential (AEP). To isolate the HEP component, an AEP subtraction procedure was applied. In the asynchronous sound condition, where the timing of heartbeats and sounds was unrelated, an AEP template was obtained for each participant by averaging EEG epochs time-locked to sound onset. This participant-specific AEP waveform was then subtracted from each trial in the synchronous sound condition. After artifact rejection, the mean numbers of remaining epochs were 935.40 for the synchronous sound condition, 1031.80 for the asynchronous sound condition, 78.63 for the synchronous omission condition, and 77.45 for the asynchronous omission condition. Although fewer epochs remained in the omission conditions than in the sound-present conditions, approximately 77–79 trials were retained on average in each omission condition, providing sufficient data for the HEP analyses.

To determine the Regions of Interest (ROI) for condition comparisons, a functional localizer analysis independent of condition differences was conducted using the non-parametric cluster-based permutation test (Maris & Oostenveld, 2007) implemented in the FieldTrip toolbox (Oostenveld et al., 2011). Because auditory-evoked activity was expected to contribute more strongly to the EEG signal after approximately 300 ms, the localizer analysis was restricted to the time window before 300 ms to minimize potential AEP contamination. We identified spatiotemporal regions in which the HEP showed a significant negative deviation from baseline (0 μV) in both the synch and asynch omission conditions. A cluster was formed when t-values exceeded the threshold (*p* < .05) at adjacent electrodes within 40 mm. The null distribution was generated using 2,000 Monte Carlo permutations (α = .05, one-tailed). The final ROI consisted of electrodes and time windows included in significant clusters for both omission conditions. This common region (E1, E2, E3, E4, E5, E6, E7, E9, E11, E51, E53, E54, E56, E57, E58, E59, E60, E61, E65; 200–300 ms) reflects stable heartbeat-locked cortical activity across auditory synchrony conditions.

A Bayesian statistical model was then constructed for the mean HEP amplitude within this ROI across five conditions (synch sound, asynch sound, synch omission, asynch omission, and rest). The synch omission condition was set as the reference category. Participants were included as a random effect (random intercept). Priors were set as *Normal* (0, 1) for fixed effects and *Student-t* (*df* = 3, μ = 0, σ = 10) for random effects. Posteriors were estimated using MCMC with 4,000 iterations.

#### 2.6.3. HCT and TET performance

Performance scores for the HCT were calculated for each trial using the formula:

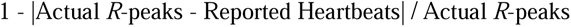

The mean score across six trials was used as the index of interoceptive accuracy.

TET performance was calculated using the formula:

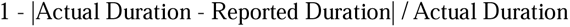

#### 2.6.4. The difference in reaction time (RT) between the experimental conditions

To assess whether sound omissions altered behavioural responses to the next auditory stimulus after sound resumption, *RT* was analysed. *RT* was calculated as the interval between the target sound presentation and the subsequent key press. Error trials (e.g., missed responses) were excluded. *RT_diff_* was calculated as the difference between the RT to the first auditory stimulus presented after an omission and the mean RT for all other regular sound-present trials, and was used as an index of interference.Trials exceeding ± 3 *SD* were excluded as outliers. Paired *t*-tests were used to compare mean *RT* and *RT_diff_* between synch and asynch conditions.

#### 2.6.5. Temporal progression of RR interval changes across trials

To assess habituation, a linear mixed model was used with Trial Number (1–15), Epoch (RR0–RR5), and their interaction as fixed effects. Participants were included as a random intercept. Posterior distributions were estimated using Hamiltonian Monte Carlo with weakly informative priors (*Normal* (0, 1)). Convergence was confirmed with *R*-hat ≦ 1 for all parameters.

2.6.6. Association with individual traits

The correlation between individual traits (BDI, STAI-Trait, BPQ, MAIA, HCT, and TET scores) and the magnitude of heart rate changes (mean RR0–RR5 rate) and HEP amplitude during omissions was analyzed. *p*-values for correlation tests were corrected using the Benjamini-Hochberg (BH) method for each condition.

## 3. Results

### 3.1. Heart rate changes following sound omissions

Before fitting the Bayesian hierarchical model, we first summarized the unstandardized RR interval changes (in ms) immediately surrounding sound omissions in each omission condition (Tables 1 and 2). For each heartbeat index (RR0-RR5), we calculated the median, mean, and standard deviation of the RR-interval change relative to the pre-omission interval (RR-1) across participants.

**Table 1.** Observed heart rate changes immediately following sound omissions (ms)

| Asynch Omission |  |  |  |  |  |  |
| --- | --- | --- | --- | --- | --- | --- |
| Statistic (ms) | RR0 | RR1 | RR2 | RR3 | RR4 | RR5 |
| Median | 2.78 | 9.57 | 5.16 | 1.8 | -0.75 | -0.58 |
| Mean | 2.54 | 9.47 | 4.87 | 0.96 | -0.44 | -0.41 |
| SD | 17.87 | 26.2 | 28.31 | 29.8 | 31.57 | 33.44 |
| Synch Omission |  |  |  |  |  |  |
| Statistic (ms) | RR0 | RR1 | RR2 | RR3 | RR4 | RR5 |
| Median | 2.33 | 12.6 | 11.54 | 8.15 | 5.27 | 5.56 |
| Mean | 2.85 | 13.76 | 12.48 | 7.82 | 3.45 | 4.49 |
| SD | 24.33 | 30.59 | 33.01 | 32.84 | 37.09 | 38.09 |

**Table 2.**
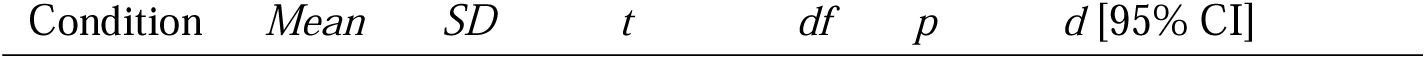
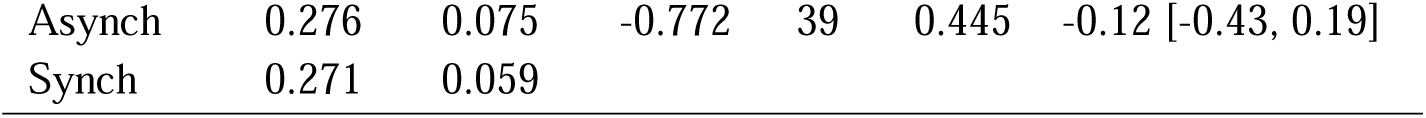
T-test statistics for the difference in mean RT between conditions.

| Condition | Mean | SD | $t$ | $df$ | $p$ | $d$ [95% CI] |
| --- | --- | --- | --- | --- | --- | --- |
| Asynch | 0.276 | 0.075 | -0.772 | 39 | 0.445 | -0.12 [-0.43, 0.19] |
| Synch | 0.271 | 0.059 |  |  |  |  |

In the asynch omission condition (Table 1), mean RR changes were small throughout the analysis window. The mean change at RR0 was close to zero (2.54 ms), and a modest RR prolongation was observed at RR1 (9.47 ms), with smaller mean increases at RR2–RR3 (4.87 ms and 0.96 ms) and values near zero at RR4–RR5 (−0.44 ms and −0.41 ms). Median values showed a similar pattern (2.78 ms at RR0, 9.57 ms at RR1, 5.16 ms at RR2, and ≤ 2 ms thereafter), indicating that, on average, asynchronous omissions induced only mild and transient heart rate deceleration.

By contrast, in the synch omission condition (Table 2), RR-interval prolongation was larger and more sustained. Although the mean change at RR0 was again close to zero (2.85 ms), mean RR changes were clearly increased at RR1 and RR2 (13.76 ms and 12.48 ms) and remained positive at RR3–RR5 (7.82 ms, 3.45 ms, and 4.49 ms, respectively). Median values followed the same trend. These descriptive results indicate a more pronounced and prolonged RR-interval prolongation following synch omissions than following asynch omissions.

A Bayesian statistical model was constructed to examine the effects of epoch (RR0–RR5) and condition on standardized RR interval fluctuations, including the main impacts of epoch and condition as well as their interaction. In this analysis, the synch omission condition was set as the reference category for comparisons against all other conditions (asynch omission, synch sound, asynch sound, and rest).

First, examining the main effects of epoch within the reference synch omission condition revealed significant positive coefficients across the RR1 to RR5 analysis window (*β* = 0.020 [0.016, 0.024] for RR1; *β* = 0.021 [0.017, 0.025] for RR2; *β* = 0.013 [0.009, 0.017] for RR3; *β* = 0.009 [0.005, 0.013] for RR4; and *β* = 0.009 [0.005, 0.013] for RR5). These results indicate a sustained prolongation of the RR interval (heart rate deceleration) starting from the first heartbeat immediately following the omission (RR1) in the synch omission condition.

Next, interaction effects between the reference condition and other conditions were evaluated. Comparison with the asynch omission condition revealed significant negative interactions at RR2 (*β* = -0.011 [-0.016, - 0.005]), RR3 (*β* = -0.008 [-0.014, -0.002]), and RR4 (*β* = -0.007 [-0.012, -0.001]). This suggests that the heart rate deceleration following an omission is significantly smaller in the asynch condition than in the synch condition during the mid-latency period (RR2–RR4).

Furthermore, marked differences were observed in conditions where auditory stimuli were actually presented compared to the omission conditions. In the synch sound condition, significant negative coefficients were confirmed from RR1 to RR3 (*β* = -0.020 [-0.025, -0.014] for RR1; *β* = -0.020 [-0.026, -0.015] for RR2; *β* = - 0.011 [-0.017, -0.006] for RR3). Similarly, the asynch sound condition showed significant negative coefficients from RR1 to RR4 (*β* = -0.018 [-0.023, -0.012] for RR1; *β* = -0.020 [-0.026, -0.015] for RR2; *β* = -0.012 [-0.017, -0.006] for RR3; *β* = -0.007 [-0.013, -0.001] for RR4). These results indicate that the early RR-interval prolongation observed after omissions was not evident during sound-present trials.

Comparison with the rest condition revealed significant negative interactions across all epochs from RR1 to RR5 (*β* = -0.019 [-0.025, -0.013] for RR1; *β* = -0.019 [-0.025, -0.013] for RR2; *β* = -0.012 [-0.017, -0.006] for RR3; *β* = -0.007 [-0.013, -0.001] for RR4; *β* = -0.007 [-0.013, -0.001] for RR5).

Collectively, these findings demonstrate that the synch omission condition induces a significantly greater degree of heart rate deceleration (RR interval prolongation) compared to sound-present and rest conditions, as well as the asynch omission condition, particularly at specific latencies. The divergence from sound-present and rest conditions indicates that the RR-interval prolongation was not explained simply by sound timing or task engagement, but was most evident when a scheduled tone was omitted in the heartbeat-synchronous context. The distribution of estimated coefficients for each variable is shown in Figure 2, and the RR interval change rates for each condition and epoch are summarized in Figure 3. Detailed parameter estimates for the model are provided in Table A1 of the Appendix.

**Figure 2.**
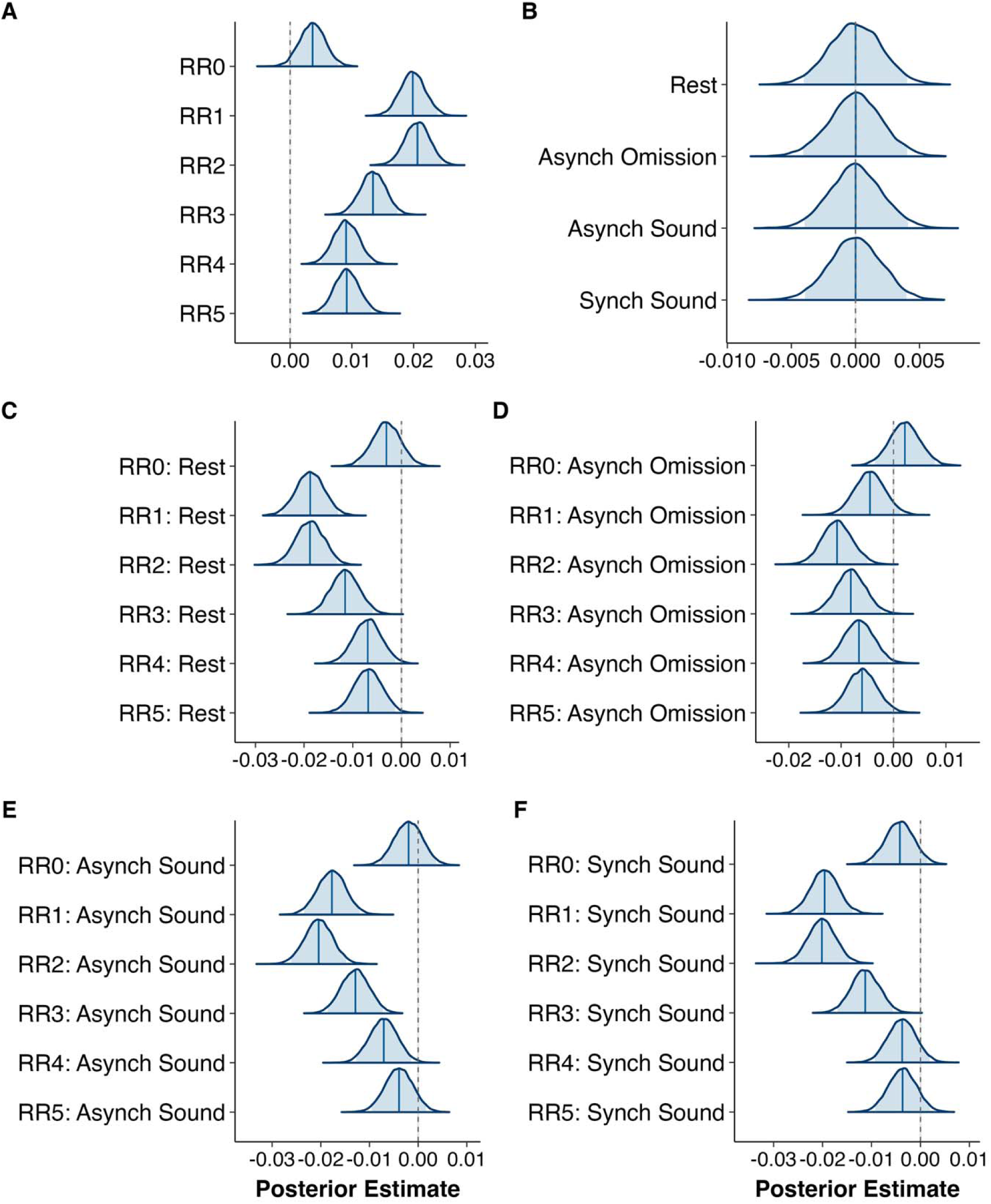
Posterior distributions of regression coefficients for fixed effects and interaction terms in the model. Across all panels, the x-axis represents the estimated values of the coefficients, and the y-axis represents the names of the indices or experimental conditions. The shaded areas in each distribution indicate the 95% Bayesian credible intervals, the vertical line located at the center of each distribution represents the posterior median, and the solid vertical line at *x* = 0 serves as the reference line indicating a coefficient of zero. The contents of each panel are as follows: (A) main effects of heartbeat epochs (RR0–RR5); (B) main effects of the four experimental conditions (rest, asynch omission, asynch sound, and synch sound). Panels (C) through (F) show the interaction terms between each experimental condition and heartbeat index: (C) rest condition; (D) asynch omission condition; (E) asynch sound condition; and (F) synch sound condition.

**Figure 3.**
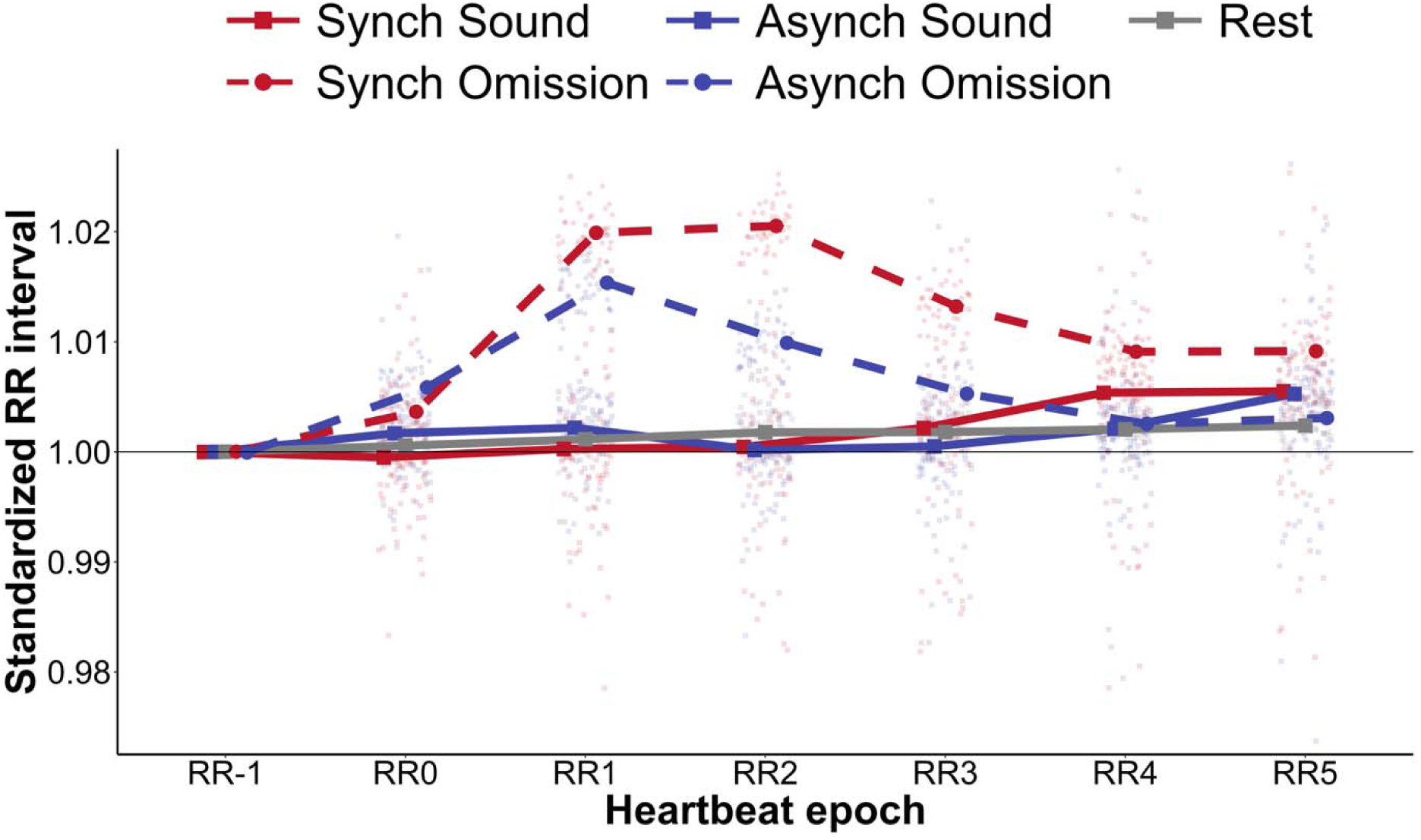
Trajectories of posterior predictive values of heart rate RR intervals across experimental conditions. This figure illustrates the changes in predicted heart rate RR intervals for each experimental condition, calculated from the posterior predictive distributions of the hierarchical Bayesian model. The x-axis represents the epochs relative to the omission or sound onset within a trial (from RR-1 to RR5), and the y-axis represents the posterior predictive values (RR). Bold lines and significant points indicate the mean values for each condition, with error bars representing the standard error of the mean. Translucent minor points in the background reflect individual posterior predictive values. Colors distinguish the primary experimental classifications: red indicates synch conditions (synch sound and synch omission), blue indicates asynch conditions (asynch sound and asynch omission), and gray indicates the rest condition. Line types and point shapes denote the status of auditory stimulation: solid lines with square points represent sound-present conditions (synch sound, asynch sound, and rest), while dashed lines with circular points represent sound-absent (omission) conditions (synch omission and asynch omission). A horizontal solid line at *y* = 1 represents the reference heart rate cycle. The results show that the omission of auditory stimuli caused a prolongation of RR intervals, with the increase being most prominent immediately following a synch omission. In contrast, no significant heart rate changes relative to the pre-stimulus period were observed immediately after actual sound presentation.

### 3.2. HEP amplitude across omission, sound-present, and rest conditions

To examine the effects of experimental conditions on the HEP, we constructed a Bayesian statistical model comparing the synch omission condition (as the reference category) against the other four conditions (asynch omission, asynch sound, rest, and synch sound). The analysis revealed significant positive coefficients for all other conditions relative to the synch omission condition: asynch omission (*β* = 0.331 [0.111, 0.550]), asynch sound (*β* = 0.455 [0.236, 0.679]), rest (*β* = 0.403 [0.177, 0.621]), and synch sound (*β* = 0.246 [0.035, 0.466]). Given that the intercept for the reference synch omission condition was negative (*β* = -0.952), the positive coefficients for the other conditions indicate a reduction in the magnitude of the negative deviation (amplitude). Thus, HEP negativity was greatest in the synch omission condition relative to all other conditions, including the synch sound condition. This pattern indicates that enhanced heartbeat-related cortical activity was most evident when a scheduled tone was omitted in the heartbeat-synchronous context, rather than during heartbeat-synchronous sound presentation itself. Figure 4A presents topographic maps averaged over the 200–300 ms time window for each condition. The distribution of the estimated coefficients for each variable is shown in Figure 4B and Table A2 (Appendix). The time-course of the EEG signals (waveforms) is illustrated in Figure 4C, and the mean HEP amplitudes averaged over the 200–300 ms window are summarized in Figure 4D.

**Figure 4.**
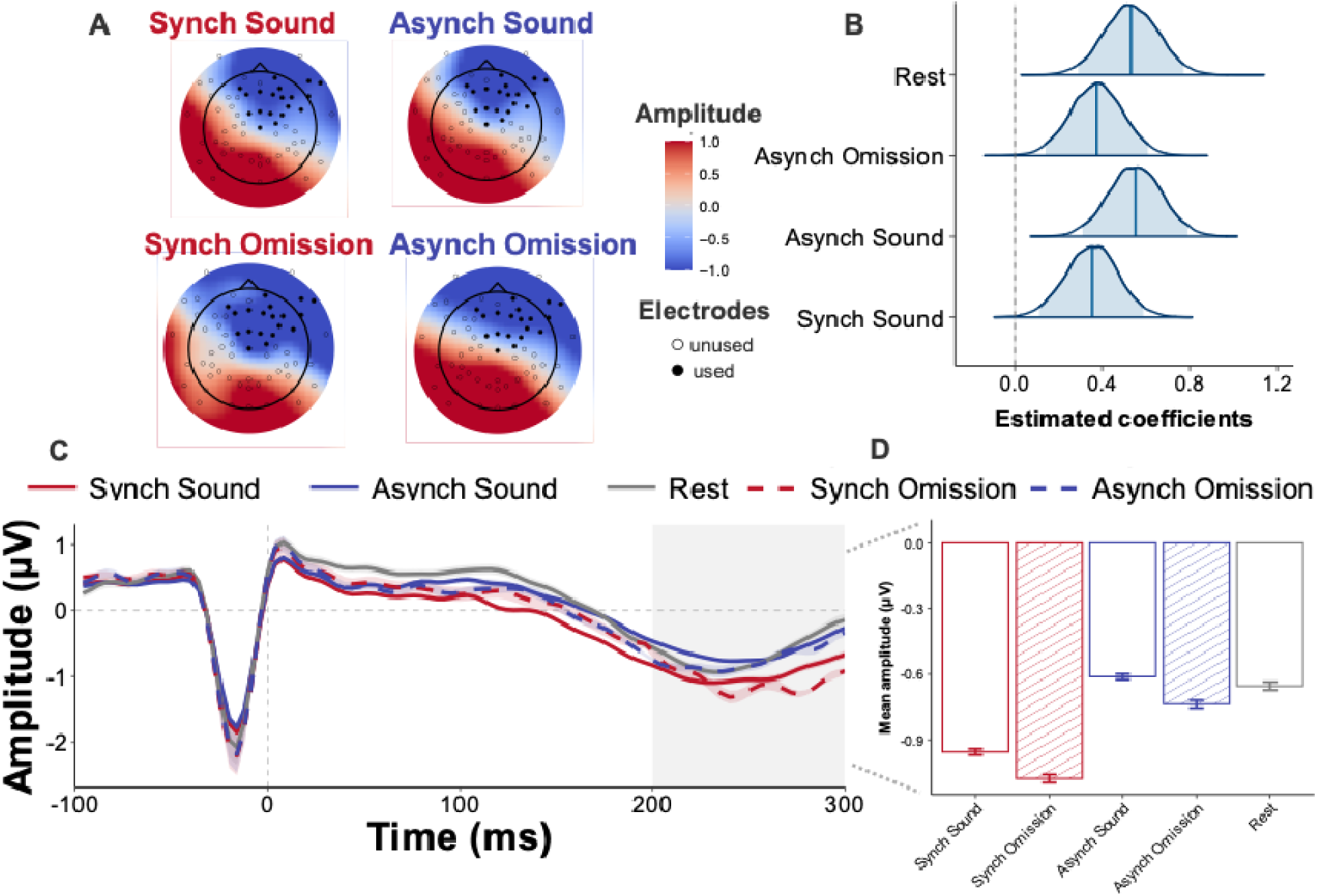
Heartbeat-evoked potentials across auditory and rest conditions. (A) Grand-average topographic maps of HEP amplitude in the 200–300 ms post-R-peak time window for the four auditory conditions: asynch omission, asynch sound, synch sound, and synch omission. The electrode cluster used for statistical analysis is indicated by black dots (E1, E2, E3, E4, E5, E6, E7, E9, E11, E51, E53, E54, E56, E57, E58, E59, E60, E61, and E65). Greater negative amplitudes were observed most prominently in the synch omission condition. (B) Posterior distributions of regression coefficients from the Bayesian model, with the synch omission condition entered as the reference category. Positive coefficients indicate less negative HEP amplitude relative to the synch omission condition. Shaded areas indicate 95% credible intervals, and vertical lines within each distribution indicate posterior medians. The vertical reference line at x = 0 indicates no difference from the synch omission condition. The posterior distributions for the asynch omission, asynch sound, rest, and synch sound conditions were shifted in the positive direction, indicating that HEP amplitudes were less negative than in the synch omission condition. (C) Grand-average HEP waveforms from the electrode cluster used for analysis. The x-axis represents time relative to the R-peak, and the y-axis represents mean amplitude (μV). Bold lines indicate condition means, and translucent shaded regions indicate standard errors of the mean. Line colours indicate timing condition (red: synch; blue: asynch), and line types indicate auditory status (solid: sound-present; dashed: omission). The vertical dashed line indicates the R-peak, the horizontal dashed line indicates zero amplitude, and the grey-shaded area indicates the 200–300 ms time window used for statistical analysis. (D) Mean HEP amplitudes in the 200–300 ms time window for the four auditory conditions. The x-axis represents condition, and the y-axis represents mean amplitude (μV). Bar colours indicate timing condition (red: synch; blue: asynch), and bar styles indicate auditory status (open bars: sound-present; striped bars: omission). Error bars indicate standard errors of the mean. The synch omission condition showed the most prominent negative HEP amplitude among the auditory conditions.

### 3.3. Behavioural results: Reaction times

Before a detailed examination of the effects of stimulus omission—the primary focus of this study—we confirmed the overall RT characteristics across conditions. A paired *t*-test conducted on the overall RTs for the synch and asynch conditions revealed no significant difference (Table 2, *t* (39) = -0.77, *p* = .445, *d* = -0.12 [-0.43, 0.19]). This result suggests that the difference in presentation conditions did not substantially influence overall response speed. Figure 5A summarizes the distribution of mean RTs for each condition.

**Figure 5.**
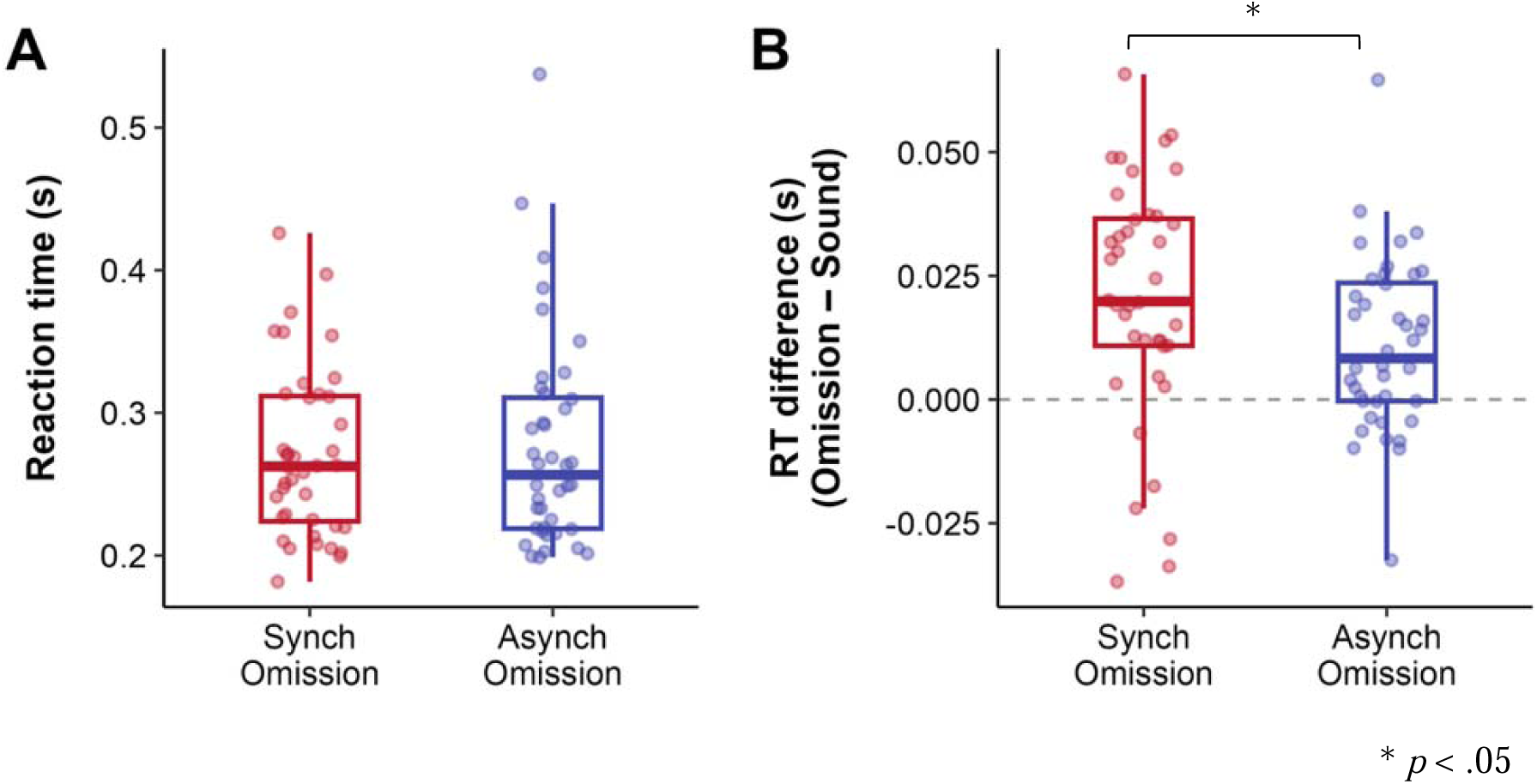
(A) Distribution of mean RT across conditions. The x-axis represents the experimental conditions (synch, asynch), and the y-axis represents the mean RT (s). The horizontal line within each box plot indicates the median, while the upper and lower boundaries represent the first and third quartiles, respectively. Individual data points for each participant are scattered around each box. The colors of the plots correspond to the experimental conditions: red for synch and blue for asynch. No significant difference in mean reaction time was observed between conditions. (B) Distribution of mean *RT_diff_* across experimental conditions. The x-axis represents the experimental conditions (synch, asynch), and the y-axis represents the difference between the RT immediately following an omission and the overall mean RT (*RT_diff_* (*s*)). The horizontal line within each box plot indicates the median, and the upper and lower boundaries represent the first and third quartiles, respectively. Individual data points for each participant are scattered around each box. The colors of the plots correspond to the experimental conditions: red for synch and blue for asynch. *RT_diff_* was significantly longer in the synch condition compared to the asynch condition.

Based on these findings, this analysis focuses on how the omission of stimuli at specific timings exerts specific interference on subsequent cognitive and behavioural processing. Specifically, to evaluate the impact of stimulus omission on subsequent key-press responses, we calculated the RT delay immediately following an omission (*RT_diff_*) relative to each participant’s mean RT in regular trials. A paired *t*-test on the *RT_diff_* for each participant revealed a significant difference between conditions (Table 3; *t* (39) = 2.60, *p* = .013, *d* = 0.41 [0.09, 0.73]). The response delay in the synch condition (0.020 ± 0.024 s) was significantly larger than that in the asynch condition (0.011 ± 0.017 s).

**Table 3.** T-test statistics for the difference in mean *RT_diff_*between conditions.

| Condition | Mean | SD | *t* | *df* | *p* | *d* [95% CI] |
| --- | --- | --- | --- | --- | --- | --- |
| Asynch | 0.011 | 0.017 | 2.598 | 39 | 0.013 | 0.41 [0.09, 0.73] |
| Synch | 0.020 | 0.024 |  |  |  |  |

This indicates that when a stimulus was presented in synchrony with the heartbeat and then omitted, it caused more substantial behavioural interference (a delay in the key-press response) in subsequent trials than in the asynch condition. Figure 5B shows the distribution of mean *RT_diff_* for each condition. These results suggest that omissions in the heartbeat-synchronous context produced a small but measurable delay in subsequent responding, consistent with transient interference in processing the first post-omission tone.

### 3.4. Temporal changes in heart rate effects across trials

To investigate whether the magnitude of the heart rate change effect fluctuated as the task progressed, we constructed Bayesian statistical models for each condition (synch omission and asynch omission), including the main impacts of epoch (RR0–RR5) and trial number, as well as their interactions. The results for both the synch and asynch conditions showed that the main effects of trial number (synch: *β* = 0.000 [0.000, 0.000]; asynch: *β* = 0.000 [0.000, 0.000]) and the coefficients for the interaction between epoch and trial number were minimal, with their 95% credible intervals mainly including zero (Appendix, Tables A3 and A4). Although the upper bounds of some interaction terms (e.g., RR4 and RR5 in the asynch condition) were slightly positive, the estimates themselves were negligible (approximately 0.001). Thus, no significant modulation of the heart rate effect by the number of trials was observed in either condition.

Figure 6 summarizes the heart rate change rates (RR-1 to RR5) for each condition across the 15 trials. While minor trial-by-trial fluctuations are visible, the synch condition consistently induced a larger heart rate deceleration (RR interval prolongation) compared to the asynch condition in almost all trials. These results suggest that the heart rate change effect in the synch condition does not attenuate but is robustly maintained throughout the task.

**Figure 6.**
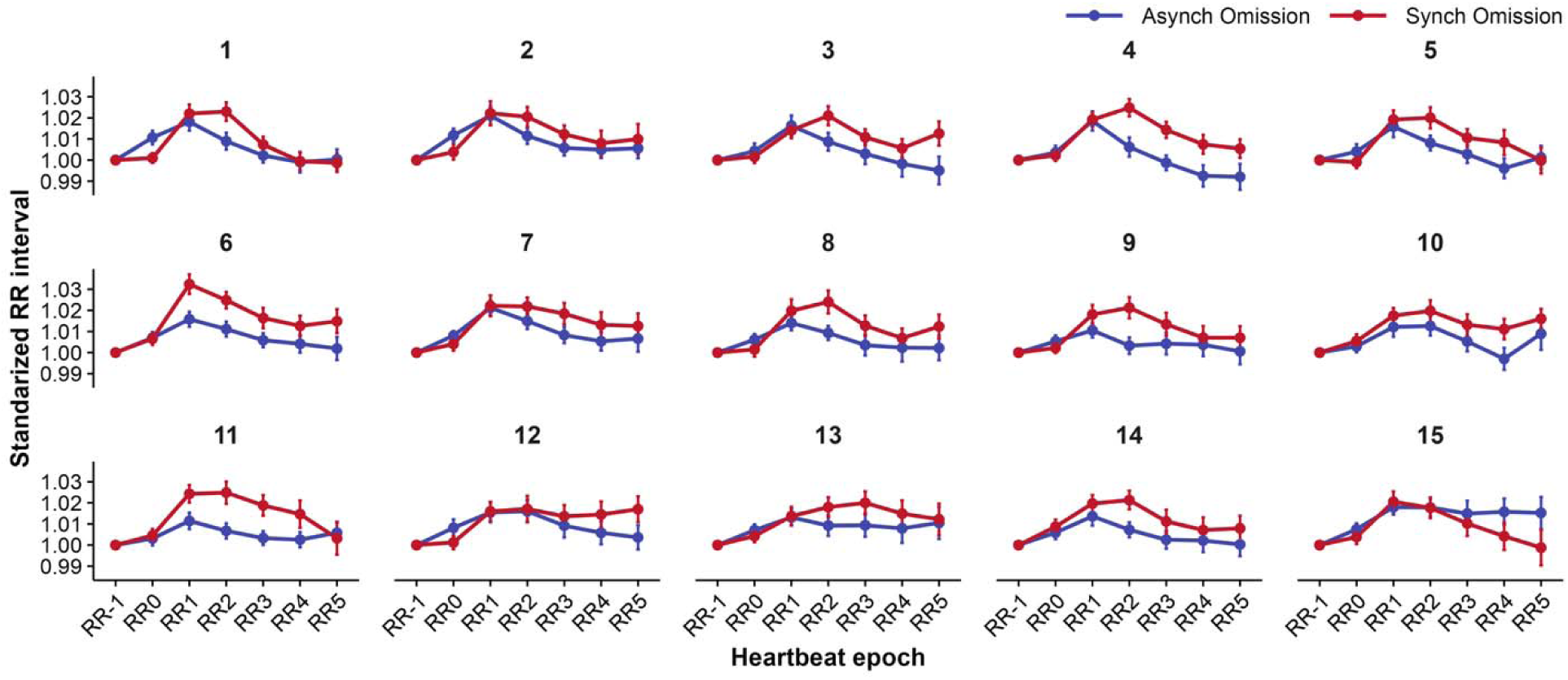
Trial-wise trajectories of standardized RR-interval changes following omissions. The x-axis represents heartbeat epochs from RR-1 to RR5, and the y-axis represents standardized RR-interval change. Each panel corresponds to a trial number, illustrating the trajectory of RR-interval changes following omissions as the task progressed. Lines and points indicate the mean standardized values for each experimental condition. In most trials, the synch condition showed greater RR-interval prolongation than the asynch condition.

### 3.5. Exploratory associations with interoceptive individual-difference measures

We explored whether omission-evoked RR-interval prolongation and HEP amplitude varied with interoceptive individual-difference measures. No clear associations were observed between these physiological responses and HCT or TET performance. Associations with interoception-related self-report measures are summarized in the Appendix (Table A5, A6). These exploratory results should be interpreted cautiously.

## 4. Discussion

The present study aimed to quantify the magnitude and duration of cardiac modulation induced by heartbeat-synchronous omissions, and to determine whether these omissions enhance heartbeat-locked cortical activity, as indexed by HEPs, relative to asynchronous omissions, sound-present trials, and rest. At the autonomic level, Bayesian hierarchical modelling showed that omissions of heartbeat-synchronous sounds induced clear and sustained RR-interval prolongation, indicating heart-rate deceleration. Specifically, coefficients for RR1–RR5 were positive in the synch-omission condition, whereas the asynch-omission condition showed significantly smaller RR-interval prolongation during RR2–RR4. Relative to both sound-present conditions and the resting baseline, synch omissions also produced larger RR prolongation across post-omission epochs. These findings support our first hypothesis by demonstrating that heartbeat-synchronous omissions evoke condition-specific autonomic modulation that is stronger and more sustained than that associated with asynchronous omissions, sound-present trials, or rest.

At the cortical level, the HEP findings showed a parallel condition-specific pattern. HEP negativity in the 200–300 ms window was greater in the synch-omission condition than in all other conditions, including asynch omissions, sound-present conditions, and rest. This effect was estimated from R-peak-locked EEG epochs falling within the interval immediately after a scheduled tone omission and before the subsequent tone presentation. Thus, the finding specifically concerns heartbeat-related cortical activity following omission of the scheduled tone in the heartbeat-synchronous context. Taken together, these findings support our second hypothesis that heartbeat-synchronous omissions selectively modulate heartbeat-related cortical responses, yielding a coupled pattern of RR-interval prolongation and enhanced HEP negativity that was most evident in the synch-omission condition.

A plausible physiological interpretation of these coupled RR and HEP effects can be framed in terms of ascending cardiac and baroreceptor-related afferent pathways and their interaction with central autonomic control circuits. Mechanical signals linked to each heartbeat are conveyed primarily via vagal and glossopharyngeal afferents to the nucleus tractus solitarius. From there, cardiac-related information ascends through brainstem and thalamic relays to insular, cingulate, and somatosensory cortices, while also interacting with medullary and vagal efferent pathways that regulate heart rate (Berntson & Khalsa, 2021; Critchley & Harrison, 2013; Ferraro et al., 2022). Within this framework, HEPs can be interpreted as a cortical signature of ascending cardiac-related input, whereas omission-evoked RR-interval prolongation may reflect autonomic output associated with heart-rate control within this broader cardio–cortical regulatory network (Kern et al., 2013; Park & Blanke, 2019).

Within this physiological framework, part of the omission-evoked cardiac response can be interpreted as a general orienting-like adjustment to auditory omission. Heart-rate deceleration is a well-characterized component of the orienting reflex to novel or unexpected non-aversive events, reflecting enhanced sensory intake and attentional orienting toward the eliciting event (Barry & Rushby, 2006; Graham & Clifton, 1966). In the present task, participants continuously monitored a repetitive auditory stream and prepared responses to tone presentations. Under this response set, even when participants were not explicitly aware of the omissions or the cardio–auditory timing rule, the absence of a scheduled tone could constitute a task-relevant deviation from the ongoing auditory sequence. The small numerical tendency toward RR-interval prolongation after asynch omissions is compatible with such a general orienting-like cardiac adjustment to auditory omission.

Beyond this general orienting-like response, the stronger and more sustained RR-interval prolongation and HEP enhancement in the synch condition suggest that auditory omissions were amplified when they occurred at a fixed latency from the participant’s own heartbeat. In the synch condition, each tone was embedded in a repeated temporal relation to cardiac afferent signalling, so the omission of a scheduled tone occurred at a time point at which cardiac-related cortical processing was also unfolding. The absence of the tone may therefore have shifted processing toward the cardiac afferent signal itself and engaged central autonomic circuits regulating heart rate. This interpretation is consistent with the combined pattern observed here: synch omissions produced multi-beat RR prolongation together with increased HEP amplitude, suggesting coordinated modulation of heartbeat-related cortical processing and autonomic cardiac control when auditory omission is tied to cardiac timing. Related work has shown that HEP amplitudes are sensitive to changes in cardiovascular state and baroreceptor input (Park & Blanke, 2019), supporting the view that cortical heartbeat responses are closely integrated with autonomic dynamics.

In this respect, the present pattern converges with and extends previous work using cardio-auditory omission paradigms. Studies employing heartbeat-locked or otherwise regular sound sequences have shown that omissions elicit larger omission-locked neural responses than omissions embedded in asynchronous or irregular sequences. When cardio–auditory regularity is present, such omissions also induce characteristic cardiac deceleration (Banellis & Cruse, 2020; Pelentritou et al., 2024, 2025; Pfeiffer & De Lucia, 2017). In line with these studies, we found that, during an active auditory attention task, heartbeat-contingent omissions produced multi-beat prolongation of RR intervals accompanied by robust HEP enhancement, while leaving overall task performance largely preserved. Comparable omission-evoked cardiac and cortical changes across passive listening, delay-judgment tasks, and the present simple detection task suggest that these responses can emerge across different task contexts, possibly reflecting relatively automatic components of cardio–cortical regulation (Critchley & Harrison, 2013; Gray et al., 2009; Park & Blanke, 2019).

At the behavioural level, omission effects were specific but limited. The RT delay immediately following omissions (*RT_diff_*) was larger in the synch than in the asynch condition, indicating a small but measurable interference effect without evidence of overall performance impairment (Figure 5). This behavioural effect should be interpreted as a transient delay in responding to the first post-omission tone, rather than as a sustained change in task performance. Trial-wise analyses of RR-interval changes further showed minimal effects of trial number and epoch × trial-number interactions, providing no clear evidence that omission-evoked cardiac responses attenuated across repeated omissions (Figure 6). Thus, heartbeat-synchronous omissions produced a brief post-omission behavioural cost, while the cardiac response to omissions recurred without clear attenuation over the course of the task. Together with the HEP findings, this pattern indicates that heartbeat-synchronous omissions can elicit autonomic cardiac modulation and heartbeat-related cortical activity even while externally directed task performance is largely preserved.

Exploratory analyses showed no clear evidence that omission-evoked RR-interval prolongation or HEP negativity varied with HCT or TET performance. This pattern suggests that the present autonomic cardiac and heartbeat-related cortical responses may reflect relatively low-level, reflex-like adjustments that are not strongly dependent on individual differences in the explicit perception or estimation of bodily signals, at least as captured by the behavioural measures used here.

This study has several limitations. First, the sample consisted of healthy young adults performing a single, simple auditory attention task. It therefore remains to be determined whether the omission-evoked cardio–cortical effects observed here generalize to other age groups, clinical populations, or tasks involving different sensory modalities and motivational contexts. Second, although we applied established preprocessing and subtraction procedures to reduce contamination of HEPs by auditory evoked activity and cardiac artefacts, these methods cannot fully exclude residual overlap. Future studies combining multiple correction approaches or, where feasible, using intracranial recordings will be important for further refining the neural interpretation of heartbeat-synchronous omission effects.

## 5. Conclusion

In summary, the present study demonstrates that omitting heartbeat-synchronous sounds during an auditory detection task produces a distinct autonomic–cortical response profile: multi-beat RR-interval prolongation and selective enhancement of HEP negativity, accompanied by only a small transient delay in responses to the first post-omission tone and no evidence of overall performance impairment. These findings extend cardio–auditory omission research from passive listening and synchrony-judgement paradigms to an active response context. They indicate that heartbeat-synchronous omissions can elicit autonomic cardiac modulation and heartbeat-related cortical activity even while auditory detection performance is largely preserved, supporting the view that heart–brain loop dynamics continue to shape physiological responses during ongoing behaviour.

## CRediT authorship contribution statement

**M.S.:** Writing – original draft, Software, Methodology, Data curation, Investigation, Funding acquisition, Formal analysis, Conceptualization.

**Y.T.:** Writing – review & editing, Methodology. **K.S.:** Writing – review & editing, Formal analysis. **Y.T.:** Writing – review & editing.

**S.U.:** Writing – review & editing, Supervision.

## Ethics approval and consent to participate

The study was approved by the Keio University Research Ethics Committee, Japan (Approval No. 240040000; approved 7 May 2024), and was conducted in accordance with the Declaration of Helsinki and relevant institutional guidelines. All participants provided written informed consent before participation, and participants’ privacy rights were observed.

## Declaration of generative AI and AI-assisted technologies in the manuscript preparation proces**s**

During the preparation of this work, the author used ChatGPT, developed by OpenAI, for language editing, wording refinement, and assistance in improving the clarity and organization of the manuscript. The author reviewed and edited the output as needed and take full responsibility for the content of the published article.

## Acknowledgement

We are grateful to all participants.

## Data availability

The datasets analyzed during the current study are not publicly available because public data sharing was not covered by the approved ethics protocol or by participants’ informed consent. The datasets contain sensitive personal information, including physiological recordings and questionnaire responses, which prevents deposition in a public repository. De-identified data may be made available from the corresponding author upon reasonable request, subject to institutional and ethical restrictions.

