## Appendix for "Autonomic and cortical responses to heartbeat-synchronous auditory omissions during an auditory attention task"

Table A1

Estimated coefficients from the Bayesian hierarchical model of omission-evoked RR-interval changes

| Parameters | Estimate | Est. error | l-95%CI | u-95%CI | Rhat | Bulk_ESS | Tail_ESS |
| --- | --- | --- | --- | --- | --- | --- | --- |
| Intercept | 1.000 | 0.001 | 0.997 | 1.003 | 1 | 2559.895 | 4705.092 |
| RR0 | 0.004 | 0.002 | 0.000 | 0.008 | 1 | 3653.011 | 6487.183 |
| RR1 | 0.020 | 0.002 | 0.016 | 0.024 | 1 | 3815.986 | 7028.309 |
| RR2 | 0.021 | 0.002 | 0.017 | 0.025 | 1 | 3833.738 | 6249.240 |
| RR3 | 0.013 | 0.002 | 0.009 | 0.017 | 1 | 3429.038 | 6290.219 |
| RR4 | 0.009 | 0.002 | 0.005 | 0.013 | 1 | 3706.572 | 6218.620 |
| RR5 | 0.009 | 0.002 | 0.005 | 0.013 | 1 | 3732.879 | 6014.956 |
| Rest | 0.000 | 0.002 | -0.004 | 0.004 | 1 | 3416.136 | 5461.237 |
| Asynch Omission | 0.000 | 0.002 | -0.004 | 0.004 | 1 | 3208.706 | 4822.777 |
| Asynch Sound | 0.000 | 0.002 | -0.004 | 0.004 | 1 | 3346.257 | 5506.749 |
| Synch Sound | 0.000 | 0.002 | -0.004 | 0.004 | 1 | 3400.684 | 5508.730 |
| RR0:Rest | -0.003 | 0.003 | -0.009 | 0.003 | 1 | 4557.373 | 6726.523 |
| RR1:Rest | -0.019 | 0.003 | -0.025 | -0.013 | 1 | 4807.333 | 7478.170 |
| RR2:Rest | -0.019 | 0.003 | -0.025 | -0.013 | 1 | 5076.001 | 6930.271 |
| RR3:Rest | -0.012 | 0.003 | -0.017 | -0.006 | 1 | 4895.377 | 6601.345 |
| RR4:Rest | -0.007 | 0.003 | -0.013 | -0.001 | 1 | 4560.271 | 7298.602 |
| RR5:Rest | -0.007 | 0.003 | -0.013 | -0.001 | 1 | 4791.254 | 7409.450 |
| RR0:Asynch Omission | 0.002 | 0.003 | -0.004 | 0.008 | 1 | 4717.283 | 6960.470 |
| RR1:Asynch Omission | -0.005 | 0.003 | -0.010 | 0.001 | 1 | 4695.962 | 6467.087 |
| RR2:Asynch Omission | -0.011 | 0.003 | -0.016 | -0.005 | 1 | 4397.587 | 7215.033 |
| RR3:Asynch Omission | -0.008 | 0.003 | -0.014 | -0.002 | 1 | 4428.210 | 7595.614 |
| RR4:Asynch Omission | -0.007 | 0.003 | -0.012 | -0.001 | 1 | 4677.789 | 6435.350 |
| RR5:Asynch Omission | -0.006 | 0.003 | -0.012 | 0.000 | 1 | 4568.118 | 6354.898 |
| RR0:Asynch Sound | -0.002 | 0.003 | -0.008 | 0.004 | 1 | 4560.568 | 7745.052 |
| RR1:Asynch Sound | -0.018 | 0.003 | -0.023 | -0.012 | 1 | 4847.232 | 7708.085 |
| RR2:Asynch Sound | -0.020 | 0.003 | -0.026 | -0.015 | 1 | 4688.817 | 6662.517 |
| RR3:Asynch Sound | -0.013 | 0.003 | -0.019 | -0.007 | 1 | 4530.967 | 7754.888 |
| RR4:Asynch Sound | -0.007 | 0.003 | -0.013 | -0.001 | 1 | 4836.358 | 7591.084 |
| RR5:Asynch Sound | -0.004 | 0.003 | -0.010 | 0.002 | 1 | 4797.129 | 7331.525 |
| RR0:Synch Sound | -0.004 | 0.003 | -0.010 | 0.001 | 1 | 4892.765 | 7394.597 |
| RR1:Synch Sound | -0.020 | 0.003 | -0.025 | -0.014 | 1 | 4878.005 | 7088.795 |
| RR2:Synch Sound | -0.020 | 0.003 | -0.026 | -0.015 | 1 | 4907.330 | 7844.000 |
| RR3:Synch Sound | -0.011 | 0.003 | -0.017 | -0.006 | 1 | 4676.822 | 7260.196 |
| RR4:Synch Sound | -0.004 | 0.003 | -0.009 | 0.002 | 1 | 4707.961 | 8029.280 |
| RR5:Synch Sound | -0.004 | 0.003 | -0.009 | 0.002 | 1 | 4804.409 | 6771.673 |

Table A2

Estimated coefficients from the Bayesian model of HEP amplitude across conditions

| Parameters | Estimate | Est. error | l-95%CI | u-95%CI | Rhat | Bulk_ESS | Tail_ESS |
| --- | --- | --- | --- | --- | --- | --- | --- |
| Intercept | -0.952 | 0.089 | -1.131 | -0.777 | 1 | 10613.493 | 8002.473 |
| Asynch Omission | 0.331 | 0.112 | 0.111 | 0.550 | 1 | 13014.158 | 9487.987 |
| Asynch Sound | 0.455 | 0.112 | 0.236 | 0.679 | 1 | 13843.753 | 10216.868 |
| Rest | 0.403 | 0.113 | 0.177 | 0.621 | 1 | 13328.477 | 9438.773 |
| Synch Sound | 0.246 | 0.111 | 0.035 | 0.466 | 1 | 13664.088 | 10136.959 |

Table A3

Estimated coefficients from the trial-number-by-RR-epoch interaction model in the synch condition

| Parameters | Estimate | Est. error | l-95%CI | u-95%CI | Rhat | Bulk_ESS | Tail_ESS |
| --- | --- | --- | --- | --- | --- | --- | --- |
| Intercept | 1.000 | 0.002 | 0.995 | 1.004 | 1 | 4374.553 | 6416.958 |
| Trial | 0.000 | 0.000 | 0.000 | 0.000 | 1 | 10592.386 | 9034.886 |
| RR0 | 0.002 | 0.003 | -0.004 | 0.007 | 1 | 11950.371 | 8980.855 |
| RR1 | 0.022 | 0.003 | 0.016 | 0.027 | 1 | 11977.576 | 9163.656 |
| RR2 | 0.024 | 0.003 | 0.018 | 0.029 | 1 | 11937.668 | 9075.831 |
| RR3 | 0.013 | 0.003 | 0.008 | 0.019 | 1 | 12178.520 | 9124.908 |
| RR4 | 0.007 | 0.003 | 0.001 | 0.013 | 1 | 12346.115 | 9944.761 |
| RR5 | 0.008 | 0.003 | 0.002 | 0.015 | 1 | 12541.062 | 10083.947 |
| RR0:Trial | 0.000 | 0.000 | 0.000 | 0.001 | 1 | 11801.173 | 10589.811 |
| RR1: Trial | 0.000 | 0.000 | -0.001 | 0.000 | 1 | 11683.941 | 10679.983 |
| RR2: Trial | 0.000 | 0.000 | -0.001 | 0.000 | 1 | 11866.899 | 10051.550 |
| RR3: Trial | 0.000 | 0.000 | -0.001 | 0.001 | 1 | 11964.732 | 9827.475 |
| RR4: Trial | 0.000 | 0.000 | 0.000 | 0.001 | 1 | 11857.928 | 10093.107 |
| RR5: Trial | 0.000 | 0.000 | -0.001 | 0.001 | 1 | 12264.828 | 9878.142 |

Table A4

Estimated coefficients from the trial-number-by-RR-epoch interaction model in the asynch condition

| Parameters | Estimate | Est. error | l-95%CI | u-95%CI | Rhat | Bulk_ESS | Tail_ESS |
| --- | --- | --- | --- | --- | --- | --- | --- |
| Intercept | 1.000 | 0.002 | 0.996 | 1.004 | 1 | 8458.820 | 8557.888 |
| Trial | 0.000 | 0.000 | 0.000 | 0.000 | 1 | 12022.923 | 10330.640 |
| RR0 | 0.007 | 0.003 | 0.001 | 0.012 | 1 | 12830.331 | 10039.283 |
| RR1 | 0.018 | 0.003 | 0.012 | 0.023 | 1 | 13703.463 | 9604.634 |
| RR2 | 0.008 | 0.003 | 0.003 | 0.014 | 1 | 12893.414 | 9986.430 |
| RR3 | 0.002 | 0.003 | -0.004 | 0.007 | 1 | 13254.386 | 10140.054 |
| RR4 | -0.002 | 0.003 | -0.008 | 0.004 | 1 | 13511.546 | 10301.807 |
| RR5 | -0.004 | 0.003 | -0.010 | 0.002 | 1 | 13668.802 | 9344.026 |
| RR0:Trial | 0.000 | 0.000 | -0.001 | 0.001 | 1 | 12102.440 | 9926.032 |
| RR1: Trial | 0.000 | 0.000 | -0.001 | 0.000 | 1 | 13471.113 | 10718.904 |
| RR2: Trial | 0.000 | 0.000 | 0.000 | 0.001 | 1 | 12696.521 | 10573.663 |
| RR3: Trial | 0.000 | 0.000 | 0.000 | 0.001 | 1 | 13195.460 | 11168.786 |
| RR4: Trial | 0.001 | 0.000 | 0.000 | 0.001 | 1 | 13029.485 | 10966.470 |
| RR5: Trial | 0.001 | 0.000 | 0.000 | 0.002 | 1 | 13192.566 | 8264.680 |

Table A5

Associations between mean post-omission RR-interval changes and interoceptive tendencies by condition

| Measure | Condition | *r* | *p* |  | Measure | Condition | | *r* | | *p* |
| --- | --- | --- | --- | --- | --- | --- | --- | --- | --- | --- |
| HCT score | Synch | 0.125 | 0.680 |  | Not-Worrying | Synch | -0.187 | | 0.680 | |
|  | Asynch | 0.056 | 0.938 |  |  | Asynch | 0.016 | | 0.938 | |
| TET score | Synch | 0.240 | 0.642 |  | Attention Regulation | Synch | 0.150 | | 0.680 | |
|  | Asynch | 0.013 | 0.938 |  |  | Asynch | 0.235 | | 0.938 | |
| BPQ | Synch | -0.236 | 0.642 |  | Emotional Awareness | Synch | 0.018 | | 0.928 | |
|  | Asynch | -0.018 | 0.938 |  |  | Asynch | -0.075 | | 0.938 | |
| Noticing | Synch | -0.122 | 0.680 |  | Self-Regulation | Synch | 0.104 | | 0.680 | |
|  | Asynch | -0.049 | 0.938 |  |  | Asynch | 0.142 | | 0.938 | |
| Not-Distracting | Synch | 0.015 | 0.928 |  | Body Listening | Synch | 0.114 | | 0.680 | |
|  | Asynch | 0.109 | 0.938 |  |  | Asynch | 0.106 | | 0.938 | |
|  |  |  |  |  | Trusting | Synch | 0.347 | | 0.393 | |
|  |  |  |  |  |  | Asynch | 0.354 | | 0.351 | |

Table A6

Associations between mean post-omission HEP amplitude and interoceptive or psychiatric tendencies by condition

| Measure | Condition | *r* | *p* |  | Measure | | Condition | | *r* | | *p* |
| --- | --- | --- | --- | --- | --- | --- | --- | --- | --- | --- | --- |
| HCT score | Synch | -0.061 | 0.899 |  | Not-Worrying | Synch | | 0.276 | | 0.556 | |
|  | Asynch | 0.019 | 0.989 |  |  | Asynch | | -0.162 | | 0.763 | |
| TET score | Synch | -0.273 | 0.556 |  | Attention Regulation | | Synch | | -0.137 | | 0.879 |
|  | Asynch | -0.013 | 0.989 |  |  |  | Asynch | | 0.165 | | 0.763 |
| BPQ | Synch | 0.021 | 0.899 |  | Emotional Awareness | | Synch | | 0.056 | | 0.899 |
|  | Asynch | -0.119 | 0.763 |  |  |  | Asynch | | 0.136 | | 0.763 |
| Noticing | Synch | 0.045 | 0.899 |  | Self-Regulation | | Synch | | -0.022 | | 0.899 |
|  | Asynch | 0.134 | 0.763 |  |  |  | Asynch | | 0.315 | | 0.364 |
| Not-Distracting | Synch | -0.076 | 0.899 |  | Body Listening | | Synch | | 0.022 | | 0.899 |
|  | Asynch | 0.002 | 0.989 |  |  |  | Asynch | | 0.309 | | 0.364 |
|  |  |  |  |  | Trusting | | Synch | | -0.237 | | 0.556 |
|  |  |  |  |  |  |  | Asynch | | 0.179 | | 0.763 |
